# Flexible IrO_x_ Neural Electrode for Mouse Vagus Nerve Stimulation

**DOI:** 10.1101/2022.10.12.511950

**Authors:** Tao Sun, Téa Tsaava, Joanne Peragine, Christine Crosfield, Maria Fernanda Lopez, Romil Modi, Rohit Sharma, Chunyan Li, Harbaljit Sohal, Eric H. Chang, Loren Rieth

## Abstract

Vagus nerve stimulation (VNS) is being actively explored as a treatment for multiple conditions as part of bioelectronic medicine research. Reliable and safe VNS in mouse models is a critical need for understanding mechanisms of these. We report on the development and evaluation of a microfabricated cuff electrode (*MouseFlex*) constructed of polyimide (PI) and with iridium oxide (IrO_x_) electrodes that is thermoformed to 86 µm ± 12 µm radius to interface the mouse cervical vagus nerve (r ≈ 50 µm). Innovative bench-top methods were used to evaluated the stimulation stability and electrochemical properties of electrodes. Our aggressive stimulation stability (Stim-Stab) test utilized 1 billion pulses at a 1000 Hz with a current density of 6.28 A/cm^2^ (1.51 mC/cm^2^/phase) to evaluate electrode lifetimes, and all electrodes remained functional. We also investigated the effects of thermoforming on their impedance, charge storage capacity (CSC), and charge injection capacity (CIC). The modest changes in electrochemical properties indicate that the thermoforming process was well tolerated. Thermoformed electrode safety and efficacy were evaluated *in-vivo* by performing acute VNS in mice and monitoring their heart and respiration rate as biomarkers. Their electrochemical properties were also measured before, during and after VNS. Bradycardia and bradypnea were reliably induced at stimulation currents of 100 to 200 µA, well below the *in-vivo* CIC of ~1250 µA (~0.5 mC/cm^2^), supporting their safety and efficacy. The electrode impedance increased and CIC decreased during *in-vivo* use, but largely reversed these changes in *in-vitro* testing after enzymatic cleaning, supporting their tolerance for surgical use.

## 1. Introduction

Electrical stimulation of and recording from peripheral nerves to treat conditions such as epilepsy [1], chronic pain [2], inflammation, and autoimmune diseases [3] are approaches known as bioelectronic medicine, and have exciting prospects to treat challenging conditions [4][5]. The human vagus nerve (AKA tenth cranial nerve) is composed of roughly 100,000 nerve fibers, and connects visceral organs to the brain [6]. A majority of the vagus is glutametergic sensory fibers, with an additional population of cholinergic efferent fibers, a large part of the parasympathetic nervous system. This makes the vagus nerve a critical component of the autonomic nervous system and its function of supporting homeostasis. Its coordination and control of multiple organ systems make it an important neuromodulation target for new treatments [7].

There are FDA-approved vagus nerve stimulation (VNS) devices to treat epilepsy and depression [8]. It is also being explored for the treatment of rheumatoid arthritis [9], hypertension [10], obesity [11], and gastrointestinal disorders [12]. Selective VNS using flexible neural electrodes has been of particular interest in translational research [13]. Polymer-based flexible thin-film neural electrodes have been fabricated using microelectromechancial systems (MEMS) approaches to interface the delicate mouse vagus nerve. The ability to tune their mechanical properties and develop architectures to decrease tethering forces and trauma, are key benefits for these electrodes. This approach also enables diverse designs, small feature sizes, cost-effective batch fabrication, and use of sophisticate materials to optimize their electrochemical and biocompatibility properties [14]. The addition of tissue engineering and drug delivery has the potential to further improved tissue integration.

Polyimide (PI) is a commonly used polymer for microfabricated flexible thin-film electrodes. However, there are few studies regarding PI electrodes specifically designed for small peripheral nerves (e.g mouse cervical vagus), and few of these report their *in-vivo* electrochemical properties in this context. Challenges to interface the small nerve (~100 µm diameter) include achieving stable and low impedances, stimulation stability, intimate contact, and the potential for robust integration suitable for chronic use.

The increased impedance and lower total (non-areal) charge injection as electrode area decreases, particularly *in-vivo*, increase the required voltage for stimulation. This can hamper the ability to stimulate without damaging the electrode or tissue. Sputtered iridium oxide (IrO_x_) has attracted considerable interest as an electrode material offering low impedance and outstanding charge storage capacity (CSC) and charge injection capacity (CIC) for both flexible and rigid electrodes. While the beneficial electrochemcial properties of IrO_x_ are well-established, the stimulation stability of IrO_x_ stacks need further optimization. We thoroughly studied the stability of an IrO_x_ stack on microfabricated flexible PI-based electrodes using a stimulation-stability (Stim-Stab) test paradigm. It included a large number of stimulation pulses (10^9^), high frequency (10^3^ Hz), and current density up to 6.28 A/cm^2^ (1.51 mC/cm^2^/phase), generating one of the harshest stimulation environments reported to date. Optical microscopy, electrochemnical impedance spectroscopy (EIS) with equivalent circuit modeling (ECM), cyclic voltametry (CV), and voltage transient (VT) measurement were performed before and after the Stim-Stab tests to comprehensively evaluate stability of IrO_x_ stack.

Several approaches have been used to achieve intimate contact with nerves, including intraneural electrode designs such as the transverse intraneural microelectrode array (TIME), longitudinal intrafascicular electrode (LIFE), and Utah (slanted) electrode arrays [15–17]. However, intraneural implantation of these technologies into the small and delicated mouse vagus nerve which is critical for homeostatis is challenging, especially for chronic studies.

Extraneural cuff electrodes have demonstrated viability for stimulation and recording from the mouse vagus nerve. Their performance varies widely, but recordings with useful signal-to-noise ratios and stimulation with modest threshold currents have been achieved. Split ring and split cylinder structures were developed to achieve the intimate interface with the ventral nerve cord of moths and sub-millimeter nerves, respectively [18][19]. Shaping planar PI-based electrodes into 3 dimensional (3D) cuffs via thermoforming has been reported as a rat vagus nerve (350 μm diameter) interface for selective stimulation and recording [20], as well as parylene-based electrodes [21]. However, electrochemical properties, stimulation stability, and efficacy of PI-based electrodes thermoformed for the small mouse vagus nerve have not been thoroughly reported upon.

Thermoformed *MouseFlex* electrodes were evaluated *in-vitro* and *in-vivo*, including the effects from thermoforming on their electrochemical properties. We also studied the safety and efficacy of acute *in-vivo* mouse VNS with concurrent measurement of their *in-vivo* electrochemical properties. Heart and respiration rate changes were quantified as biomarkers for stimulation efficacy. Furthermore, electrochemical measurements are performed before, during and after the electrode placement procedure on the mouse cervical vagus nerve, to enable the effects of surgical handling and the *in-vivo* environment to be carefully characterized. The reversable and comparable electrochamical properties measured before and after the mouse VNS demonstrated the electrode robustness and tolarance for surgical handling and use. Importantly, the *MouseFlex* electrode can also be scaled and adapted to interface a broad range of targets in the central, peripheral, and autonomic nervous systems, making it a platform technology for neuroscience research.

## 2. Materials and methods

An exploded view of the *MouseFlex* electrodes is presented in Fig. 1a. They comprise 1) a bottom PI insulating layer (7 µm), 2) Ti/Pt/Au (100/250/425 nm) pad/trace metallization, 3) Ti/Ir/IrO_x_ (15/30/450 nm) electrode sites, and 4) the top PI insulating layer (7 µm).

**Fig. 1.**
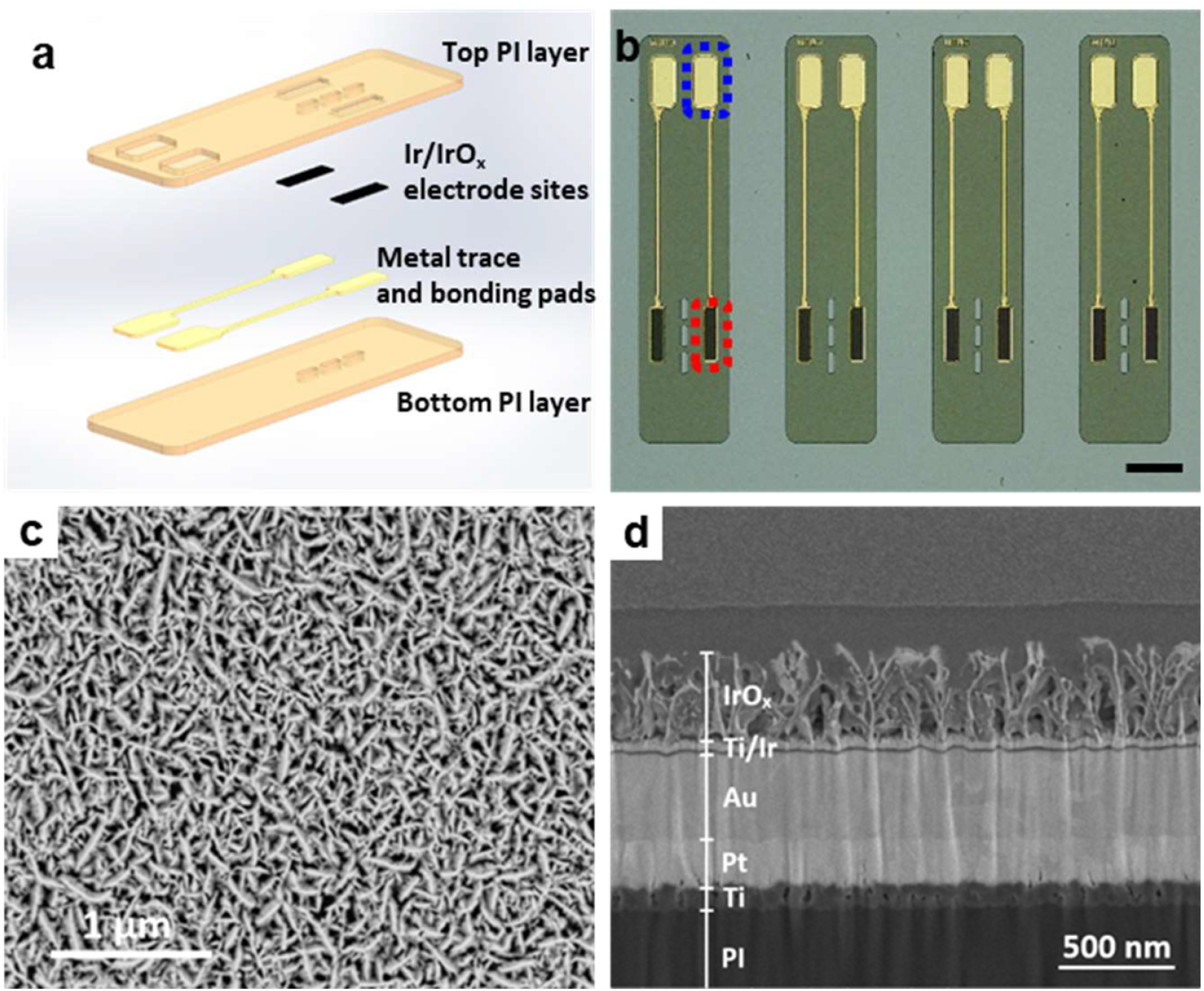
Characterization of the *MouseFlex* electrode. a) the exploded view of the *MouseFlex* electrode to reveal its structure, b) optical image of the micro-fabricated *MouseFlex* electrodes on Si wafer (the gold bonding pad is labeled by the blue dot box, and the black IrO_x_ on the electrode site is highlighted by the red dot box), scale bar = 1 mm, c) the surface morphology of the IrO_x_ on the electrode site, d) the cross-sectional morphology of the multilayered structure on the electrode site.

### 2.1 Microfabrication process

The *MouseFlex* microfabrication process is briefly outlined, and a schematic process flow is presented in Fig. S1 (supplementary materials). Electrodes were fabricated on 100-mm silicon (Si) wafers (Hoya Corporation, Tokyo, Japan), which were cleaned and rendered hydrophobic with buffered oxide etch (BOE), then rinsed and dried in a spin-rinse-dryer (SRD) using deionized (DI) water and N_2_, respectively. The bottom PI layer was then spin-coated, and soft baked at 130°C for 90 s, followed by curing at 300°C for 1 hour in N_2_, yielding 7 µm thick cured films. An O_2_ reactive ion etch (RIE) was performed at 100 W for 60 s to enhance trace metal adhesion [22]. Next, a 10-µm-thick photoresist layer of AZ9260 (MicroChemicals GmhH, Ulm, Germany) was spin-coated, baked, and processed as part of the lift-off photolithography to pattern the pad/trace metallization. A Ti/Pt/Au (100/250/425 nm) stack was then magnetron sputter deposited (Discovery 635, Denton vacuum LLC, Moorestown, NJ). Liftoff was then performed using acetone with ultrasonic agitation, followed by rinsing in isopropyl alcohol (IPA) and an SRD process. Subsequently, a second 10-µm-thick AZ9260 layer was processed to define the Ti/Ir/IrO_x_ (15/30/450 nm) electrode stack. The IrO_x_ layer was reactively sputter deposited from a metallic Ir target in an O_2_ (100 sccm) and Ar (100 sccm) plasma (TMV SS-40C-IV, TM Vacuum, Cinnaminson, NJ). The IrO_x_ film was deposited using 50 Watt from a pulsed-DC power supply, and liftoff was then performed. The top PI layer was then spin-coated and cured at 300°C for 1 h in nitrogen atmosphere, yielding a 7-µm thick insulating layer. To open the bond pads, electrode sites, perforations, and define the electrode outline, a double layer (20 µm thick) of AZ9260 was spin-coated and patterned by photolithogrphy to serve as a soft mask. An O_2_ RIE process was then performed for 51 minutes using He backside cooling in an Oxford Plasmalab 100 (Bristol, UK) to etch the exposed PI. The metallized pads and electrodes serve as etch stops for these structures while the full 14 µm of PI was etched to define the electrode outlines and perforations. Subsequently, the photoresist soft mask was removed by acetone. The geometrical surface area of each exposed IrO_x_ electrode site is 0.001274 cm^2^. The microfabricated *MouseFlex* electrodes were manually released by a scapel or tweezers.

### 2.2 Surface characterization

Optical images of *MouseFlex* electrodes were recorded by a digital microscope (VHX-7000, Keyence, Itasca, IL). The surface morphology and cross-sectional morphology of IrO_x_ electrode sites were observed using scanning electron microscopy (SEM, Apreo™, Thermo Fisher Scientific, Hillsboro, OR) at a primary beam energy of 20 kV. Dual beam-Focused Ion Beam (db-FIB) images were acquired using a Helios Nanolab 650 (Thermo Fisher Scientific, Hillsboro, OR) equipped with a 30 kV Ga beam for milling and 20 kV electron beam for imaging. The IrO_x_ porosity was quantified using ImageJ (NIH, Bethesda, MD) from both plan and cross-sectional views.

### 2.3 Electrode integration

*MouseFlex* electrodes were integrated with polytetrafluoroethylene (PTFE) insulated Pt-Ir lead wires (Ø=150 µm, Sandvik, Palm Coast, FL) for both bench studies and acute *in-vivo* VNS. Specifically, both ends of the Pt wires were mechanically de-insulated, and one end was soldered to the *MouseFlex* bond pads. A fine tipped soldering iron (NAS 2-Tool Nano, JBC Tools Inc, St. Louis, MO) and a SAC 305 lead-free solder paste with water-soluble flux (Indium Corp., Clinton, NY) were utilized. The unmelted solder paste and water-soluble flux were then removed by thoroughly rinsing and irrigation of the electrode and solder joint with warm deionized (DI) water and carefully air dried. The solder joints were mechanically reinforced by applying a small amount of medical grade epoxy (Loctite M 31 CL, Rockhill, CT), and cured at 60 °C for 45 min in an oven (Heratherm™, Thermo Fisher Scientific, Waltham, MA). The solder joints were then encapsulated with NuSiI MED 4211 medical grade silicone (Nusil Technology LLC, Carpinteria, CA) degassed in a centrifuge to electrically insulate and mechanically support them.

#### 2.3.1 Electrochemical impedance spectroscopy (EIS)

Impedance spectra of *MouseFlex* electrodes were measured a using 3-electrode configuration with one *MouseFlex* channel as the working electrode, an uninsulated Pt wire as the counter electrode. A Ag|AgCl reference electrode (Gamry Instruments, Warminster, PA) was used for bench measurements, and a Ag wire treated to form a Ag|AgCl reference for *in-vivo* measurements. *In-vitro* impedance spectra were measured in 1× phosphate buffered saline (PBS) at a *p*H of 7.4, using a Reference 600+ potentiostat (Gamry Instruments, Warminster, PA). The configuration for *in-vivo* measurements is described in section 2.6. Potentiostatic spectra were measured from 1 to 10^5^ Hz with 10 points per decade and 3 sweeps per point using a 25 mV_rms_ signal. All benchtop EIS measurements were performed in a Faraday cage. The retrieved thermoformed *MouseFlex* electrodes were soaked in ENZOL® (Advanced Sterilization Products, Irvine, CA) overnight to remove proteins or organic matter on electrodes, followed by a gentle but thorough rinsing in DI water and careful drying.

#### 2.3.2 Cyclic voltammetry (CV) measurement

The CV measurements used the same configurations as EIS, and are used to evaluate electrochemical reactions at the electrode interface and quantify their charge storage capacity (CSC, mC/cm^2^). A sweep rate of 50 mV/s was employed between the electrolysis potential (water window) limits of −0.6 and 0.8 V, with the last of 3 sweeps used to calculate their CSC. The CSC was quantified by integrating the cathodic current density across the potential range. Measurements were performed before, during, and after surgical placement.

#### 2.3.3 Voltage transient (VT) measurement

The charge injection capacity (CIC) of *MouseFlex* electrodes was determined by the voltage transient (VT) waveforms from the cathodic stimulation current pulses. They were characterized with symmetric, charge-balanced, cathodic leading, biphasic pulses with pulse-widths of 200 µs, 300 µs and 400 µs per phase, with a 40 µs interphase, at a pulse rate of 10^3^ Hz. The monopolar stimulation pulses were delivered to electrodes with an uninsulated Pt wire as the ground/counter, sourced from a neural stimulator (STG4008, Multi Channel Systems GmbH, Germany).

The maximum cathodic potential (*E*_mc_) was calculated by subtracting the access voltage (*V*_ac_), associated with the Ohmic resistance from the solution, from the maximum negative voltage in the transient (Fig. S1, supplementary materials). CIC was determined by the current amplitude and pulse width, when the *E*_mc_ reaches the water window (−0.6V) as verified by CV.

### 2.4 Stimulation-stability (Stim-Stab) test

The electrical stimulation stability of the IrO_x_ electrodes was determined by a stimulation-stability (Stim-Stab) test at room temperature (n=6). Briefly, the IrO_x_ electrode sites of *MouseFlex* electrodes were soaked in 1×PBS with a *p*H of 7.4 in small glass vials. One channel of the biopolar *MouseFlex* electrode was stimulated using a Pt wire counter/ground electrode. The unstimulated channel was retained as a control. A charge-balanced, biphasic, cathodic-leading, symmetrical stimulation paradigm with 1 billion pulses at a frequency of 10^3^ Hz, lasting 277.78 hours, was delivered to stimulated channels, using the neural stimulator. The pulse-width per phase, interphase, and interpulse period were 240 µs, 40 µs and 480 µs, respectively, using a stimulation current of 8 mA (1.51 mC/cm^2^/phase), resulting in a total dose of 1924 C. Voltage transient across a 1 kΩ resistor was monitored using an oscilloscope (DSOX2004A, Keysight Technologies, Santa Rosa, CA) and recorded with the NI DAQ to verify consistent output. The voltage waveforms from stimulated electrodes and the monitor resistor were regularly recorded using a data acquisition system (NI-9221 DAQ, National Instruments, Austin, TX), and custom MATLB (2016a, Mathworks, Natick, MA) code. Voltage waveform snippets that included 2 or 3 pulses were collected ever 250,000 pulses (250 s).

### 2.5 Thermoforming *MouseFlex* electrodes

The 2D *MouseFlex* electrodes (n=12) were shaped into 3D cuff structures, by thermoforming around 127-µm-diameter tungsten rod as a mandrel, which is close to the mouse vagus nerve diameter. The mandrel was oriented perpendicularly to the *MouseFlex* electrode and aligned with the middle of the electrode. The distal aspect of the electrode was then folded around the mandrel with fine forceps.

Heated air (260 °C setpoint) from a hot air rework station (WHA900, Weller®, Switzerland) through a fine nozzle was applied perpendicularly to the wrapped electrode for 3 mins (Fig. S2a). The process was repeated horizontally for another 3 mins (Fig. S2b), resulting in a thermoformed cuff shape for the *MouseFlex* electrode.

### 2.6 Acute mouse vagus nerve stimulation (VNS)

The protocols for *in-vivo* study regarding stimulation safety and efficacy of *MouseFlex* electrodes were approved by the Institutional Animal Care and Use Committee (IACUC) at the Feinstein Institutes for Medical Research (Manhasset, NY, USA). The Feinstein Institutes follow the National Institute of Health (NIH) guidelines for the ethical treatment of animals. The *in-vivo* electrochemical properties of *MouseFlex* electrodes were evaluated using EIS, CV (CSC), and VT (CIC) measurements for electrodes placed on the mouse cervical vagus nerve at body temperature (37 °C). EIS and CV were collected in a 3-electrode configuration using a Ag wire processed to form a Ag|AgCl reference electrode. Stimulation efficacy and VT measurements were performed concurrently by measuring changes in heart and respiration rates, and recording the voltage waveforms from the stimulation. The *in-vivo* CIC was determined from VT measurements using the previously outlined method. We monitored changes in heart and respiratory rates as the primary biomarkers. The slowed heart (bradycardia) and respiration (bradypnea or apnea) rates serve as metrics for VNS efficacy.

#### 2.6.1 Electrode placement surgery

Electrodes were placed on the left cervical vagus nerve of 10 to 12-week-old male C57BL/6J mice (n=8, JAX® Mice, Bar Harbor, Maine). The mice were anesthetized using isoflurane (3.0% induction and 1.5% maintenance). While in the supine position, a midline cervical incision was made on the neck and the left vagus nerve was isolated from the carotid artery as previously described [23]. The cervical vagus nerve was de-sheathed by removing the thin connective tissue surrounding the nerve using fine surgical forceps under magnification.

The thermoformed *MouseFlex* electrode was passed under the vagus nerve, and retracted to ensconce the nerve in the cuff and make close contact with electrode sites. A silver wire (Ø ≈ 0.2 mm) coated by AgCl reference electrode was placed between right salivary gland and the skin, while a de-insulated Pt wire counter electrode was placed between left salivary gland and the skin.

#### 2.6.2 Vagus nerve stimulation (VNS) and VT measurement

The efficacy of *MouseFlex* electrodes was evaluated by performing VNS with a range of stimulation currents and measuring changes in heart rate (∆HR) and respiration rates (∆RR), which are established biomarkers of vagus nerve engagement. VTs were also recorded during the stimulation to measure the CIC *in-vivo*. VNS was delivered at 30 Hz (pulses per second) for 7 s with cathodic-leading biphasic pulses using a pulse width of 260 µs, interphase delay of 40 µs, and current from 100 to typically 1400 µA, in 100 µA increments. VNS was delivered to both channel 1 and 2 in random order with a rest period > 1 min between stimulation trains. The order of channel stimulation was randomized to control for potential order effects. Electrocardiogram (ECG) data for HR and electrophysiological recording for respiration rates were acquired at 32 kHz using a Plexon data acquisition system (Omniplex, Plexon Inc., Dallas, TX). Data was analyzed offline using Spike2 software (Cambridge Electronics Design Limited, Cambridge, England) to quantify heart and respiration rates. The *in-vivo* VT were acquired using an NI-9221 DAQ, and recorded and analyzed with custom MATLAB code. For comparison with the *in-vivo* CIC calculated from the VNS, *in-vitro* VT was performed to obtain *in-vitro* CIC using the same parameters (frequency: 30Hz; pulse width: 260 µs; interface delay: 40 µs; temperature: 37 °C) before the electrode placement on the vagus nerve.

### 2.7 Statistical analysis

All results are present as means with the standard deviation and statistical analysis (Prism, 2020, GraphPad, San Diego, CA). Multiple comparisons were analyzed using one-way ANOVA followed by Bonferroni correction post hoc test, and *p* values <0.05 were considered statistically significant. Each channel of a *MouseFlex* electrode used in EIS, CV, and VT measurement and for Stim-Stab and *in-vivo* testing is considered as an independent sample in the statistical analysis.

## 3. Results

A representative micrograph of microfabricated *MouseFlex* electrodes on a Si wafer is presented in Fig. 1b. The relatively small area of the electrodes and compatibility with leads integrated by soldering, as well as conductive adhesives and wire bonding (not shown), yield > 100 devices per 100-mm wafer. The black area highlighted in the red dot box is the IrO_x_ electrode site, while the traces and bond pads are the reflective gold-toned structures. The transparent yellowish film is the PI used as insulation and forms the electrode architecture. The higher magnification top-view SEM image in Fig. 1c presents the surface morphology IrO_x_ layer, showing the porous dendritic structure of the film [24]. The porosity of the IrO_x_ layer from the plan view is estimated to be 25.15 ± 1.88% by ImageJ. The cross-section morphology observed from focus ion beam (FIB) processing and SEM imaging of the electrode sites reveals the multilayered structure, and a dendritic morphology with nano-sized pores for the IrO_x_ top layer (Fig. 1d). Note that a Pt “strap” was deposited in the db-FIB on the IrO_x_ electrode surface to preserve the surface morphology during cross section preparation. The rippled “curtaining” of the IrO_x_ resulted from FIB miling of the porous IrO_x_ film. The observed Ti layer was used to increase adhesion of the trace metallization and the PI substrate. This was considered effective due to the absence of observed delamination in the structures studied. The Pt, Au, and metallic Ir layers had a dense microstructure, while the IrO_x_ layer has a very high degree of porosity estimated to be 33.02 ± 2.25%. The high porosity of the IrO_x_ layer is intentional, as the process conditions yielding this morphology achieve relatively low impedance, high CSC and CIC values based on past optimizations [25–27].

The electrical stimulation stability of the electrodes was investigated using an aggressive stimulation (Stim-Stab) protocol with a large number (10^9^) of pulses and a high charge density of 1.51 mC/cm^2^/phase. One electrode site of each *MouseFlex* electrode was stimulated, while the other served as an unstimulated control to evaluate the effects of soaking time in PBS. Fig. 2a presents Bode plots of unstimulated channels before and after the Stim-Stab was performed. No statistically significant difference was observed except for a small decrease (19.83%~22.58%) from 40 Hz to 100 Hz (*p* < 0.05). Similarly, a small but statistically significant decrease (less negative) in phase angle was observed over the range of 1.5 Hz to 6.3 Hz (*p* < 0.05). Note that no significant difference was observed between unstimulated and stimulated electrodes prior to the test (*p* > 0.05), as shown in Fig 2a and b.

**Fig. 2.**
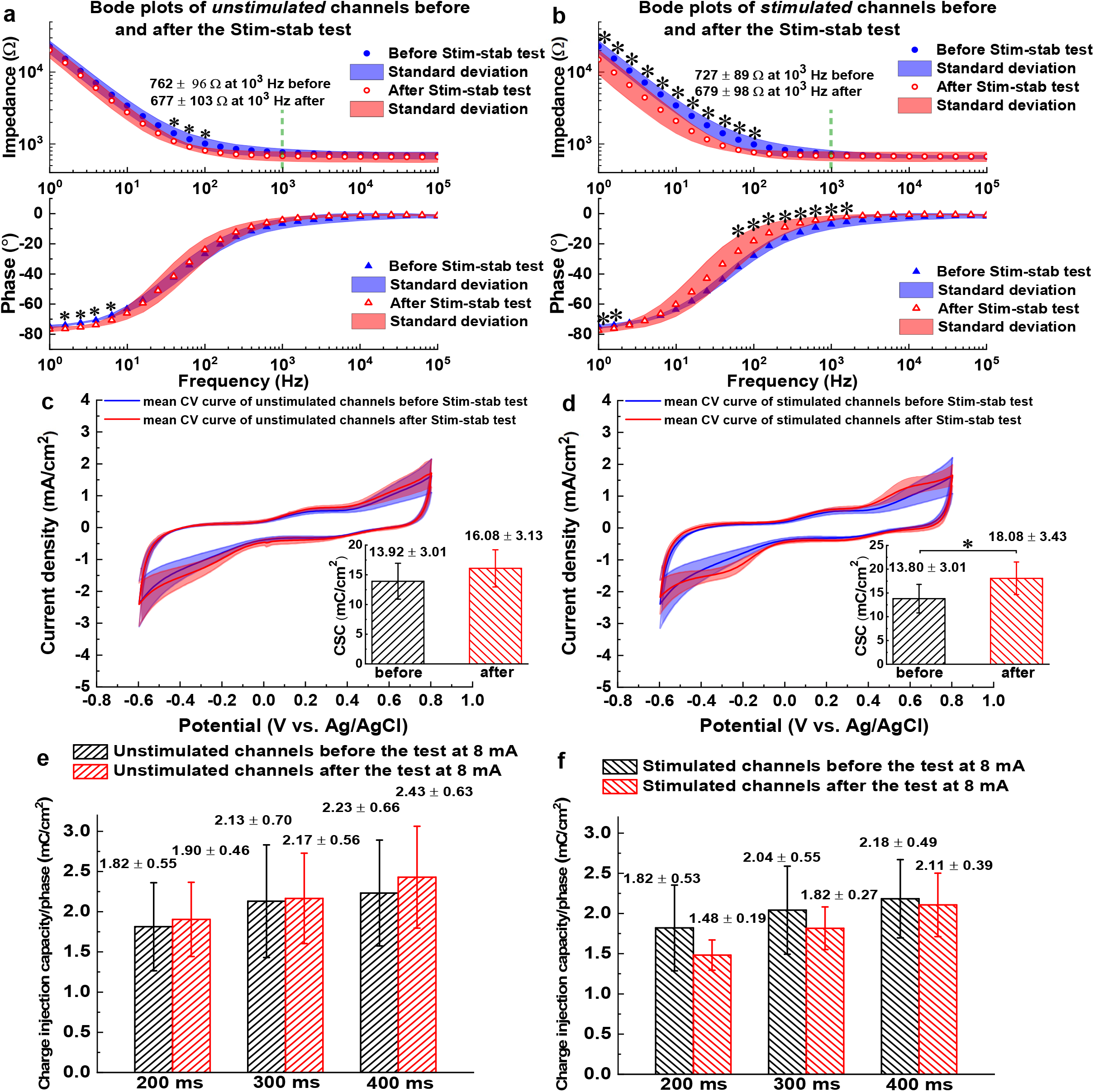
Electrochemical properties of unstimulated and stimulated channels before and after the Stim-Stab test. a) Bode plots of unstimulated channels before and after the Stim-Stab test at 8 mA, b) Bode plots of stimulated channels before and after the Stim-Stab test at 8 mA, c) the mean CV curves of unstimulated channels before and after the Stim-Stab test at 8 mA, the shaded areas represent the standard deviation, the inset figure presents CSC values of unstimulated channels before and after the Stim-Stab test, d) the mean CV curves of stimulated channels before and after the Stim-Stab test at 8 mA, the shaded areas represent the standard deviation, the inset figure shows CSC values of stimulated channels before and after the Stim-Stab test, e) CIC of unstimulated channels at different pulse width before and after the Stim-Stab test, f) CIC of stimulated channels at different pulse width before and after the Stim-Stab test.

Electrodes that were stimulated using the Stim-Stab protocol, experienced a statistically significant and larger *decrease* (22.79%~34.77%) in impedance for frequencis from 1 Hz to 100 Hz (*p* < 0.05, Fig. 2b). This represents the low-frequency region dominated by the capacitive interfacial impedance of the electrode. No statistical difference was observed in impedance at higher frequencies (*p* > 0.05), the region of the impedance spectra dominated by the access resistance. These results suggest the electrode can retain desirable electrochemical properties after aggressive Stim-Stab testing. Significant changes in phase angle were observed at low frequencies (1.0 to 1.15 Hz), where the phase angle slightly decreased as well as from 60 to 1578 Hz (*p* < 0.05) where it increased by a relatively large amount. These changes are consistent with the onset of resistive character for the electrode happening at lower frequencies compared to before stimulation of the electrode, which is in turn consistent with electrochemical activation of the IrO_x_ [28].

CV plots and the associated CSC values of unstimulated (control) and stimulated electrodes were also compared, before and after stimulation. No statistical difference in CV or CSC was observed between electrodes prior to stimulation (*p* > 0.05). CV data from unstimulated electrodes are presented in (Fig. 2c). Strong redox peaks are not present in the mean (n=6) CV curves before and after soaking the unstimulated electrodes and are consistent amongst the samples. There was a similar lack of statistical difference (*p* > 0.05) for their CSC values.

In contrast, stimulated electrodes, demonstrate two modest sized peaks centered at around 0.6 V and −0.25V (Fig. 2d). They likely result from reversible oxidation and reduction between Ir^3+^ and Ir^4+^ [27]. In addition, the CSC value of stimulated channels significantly *increased* from 13.80 ± 3.01 mC/cm^2^ to 18.08 ± 3.43 mC/cm^2^ (*p* < 0.05). However, there was no significant change (*p* > 0.05) in the CIC for stimulated or unstimulated electrodes as can be observed in Fig. 2f and 2e, respectively. The increase in CSC in conjunction with the noted decreases in impedance suggest that electrode activation might be occurring, but if so does not have a significant impact on the CIC [28]. The mechanism of stimulation induced electrode activation on CIC are not well explored, making this effect difficult to evaluate in comparison to other studies. The lack of electrode degradation, and slight improvement in electrochemical metrics despite the large stimulation doses (> 1900 C) support their outstanding resilience.

Voltage transient (VT) waveforms, *E_mc_* and *E_ac_* values from Stim-Stab, optical micrographs, and equivalent circuit models based on fits to EIS datasets for representative stimulated channels were used to further quantify and analyse degradation. Stimulated channels were compared to the unstimulated control electrode on the device (Fig. 3). The voltage waveforms are the time-domain voltages applied by the stimulator to drive the stimulation pulses. Notable voltage increases or waveform distortions imply changes, degradation, or failure of the electrode, due to changes in surface area or electrochemical properties. Electrode degradation can therefore result in increased voltages and electrical polarization. Data from electrode 1 channel 2 (denoted as E1C2) and E2C2 were selected, because they have the most significant damage observed by optical microscopy for the electrode along with relatively large fluctuation in *E*_ac_.

**Fig. 3.**
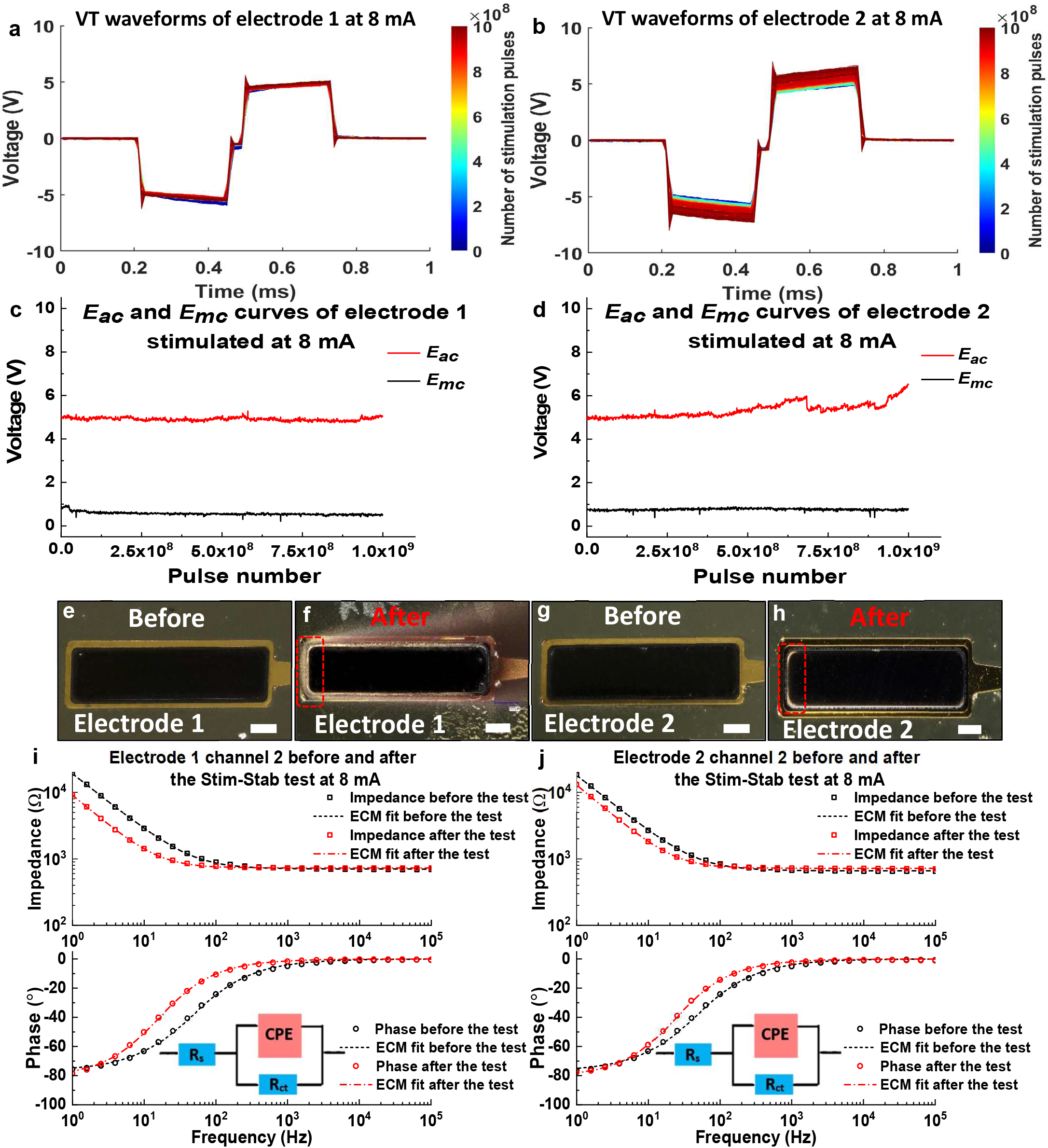
VT waveforms, *E*_ac_ and *E*_mc_ curves, optical images and ECM fitting of stimulated channels before and after the Stim-Stab test. a) VT waveforms of electrode 1 (E1C2) at 8 mA, b) VT waveforms of electrode 2 (E2C2) at 8 mA, c) *E_ac_* and *E_mc_* curves of electrode 1 stimulated at 8 mA, d) *E_ac_* and *E_mc_* curves of electrode 2 at 8 mA, e) optical image of electrode 1 before the Stim-Stab test, f) optical image of electrode 1 after the Stim-Stab test, and the red dot box highlights the typical damage area on the electrode sites, g) optical image of electrode 2 before the Stim-Stab test, h) optical image of electrode 2 after the Stim-Stab test, and the red dot box highlights the typical damage area on the electrode sites, i) Bode plots of electrode 1 (E1C2) before and after the Stim stab test, and equivalent circuit model (ECM) fit for the Bode plots, j) Bode plots of electrode 2 (E2C2) before and after the Stim stab test, and ECM fit for the Bode plots, scale bars = 100 µm.

Fig. 3a and b present the VT waveforms of, E1C2 and E2C2 as the function of the stimulation cycle, respectively. The VT from the 1 kΩ monitoring resistor had negligible variation (<2%) indicating stable output (Fig. S3). The VT measurements from electrode E1C2, are representative, and demonstrate relative stability compared to E2C2 longitudinally (Fig. 3a). Longitudinal waveform changes can be in terms of *E_ac_* values, which dominates the observed changes in voltage waveforms compared to *E_mc_*. For E1C2, the maximum increase of *E_ac_* was only 4.85% (Fig. 3c) compared to 28.4% change for E2C2 (Fig. 3d), which is the largest increase. As the mean value of the maximum increase in *E_ac_* is merely 3.66 ± 1.04% for other samples, the large fluctuation in *E*_ac_ of E2C2 suggests some degree of electrode degradation occurred.

Optical microscopy is used to evaluate electrode damage resulting from the Stim-Stab test. No damage was observed for unstimulated control electrodes after soaking (Fig. S4). Although the black-toned IrO_x_ layer remained intact for most of electrode area for the stimulated channel E1C2, a slight discoloration that potentially results from electrode degradation was noted at two corners and edges of the electrode (Fig. 3e and f, highlighted in red dot box). This potentially damaged area was quantified by ImageJ represents 4.30 ± 0.27% of the electrode area. An additional gold-toned discoloration was observed surrounding the electrode. It remained despite rinsing in deionized water suggesting a process internal to the PI encapsulation. The discoloration was opaque using transmitted light microscopy (Fig. S5) suggesting material transport at the interface of PI layers or within them.

Changes in electrochemical properties of the electrodes were quantified and evaluated by equivalent circuit modelling (ECM) to fit the Bode plots of the stimulated channel. ECM based on the classic Randles circuit [30]–[32], fits reasonably well for the Bode plots. Electrodes with degradation exhibit distinctively different parameters compared to undegraded and control. For the stimulated channel E1C2 before and after Stim-Stab test, the minor damage observed optically does not correlate with degradation of the electrochemical properties. In Fig. 3i, the impedance of E1C2 is actually lower for frequencies < 100 Hz after stimulation, which is consistent with electrode activation effects. The ECM consists of a constant phase element (CPE) shunted by a charge transfer resistance *R_ct_*, together in series with the solution resistance *R*_s_. The ECM parameters to best fit the Bode plots are presented in Table 1. One notable trend is the decrease in R*_ct_* observed for both unstimulated and stimulated electrodes. This consistency suggests the decreased charge transfer resistance is intrinsic to the soaking process of these electrodes.

**Table 1.** Parameters of the ECM for *MouseFlex* electrodes before and after the Stim-Stab test.

|  | Before Stim-Stab test |  |  |  | After Stim-Stab test |  |  |  |
| --- | --- | --- | --- | --- | --- | --- | --- | --- |
| Unstimulated channels | $R_s$ ($\Omega$) | $R_{ct}$ ($G\Omega$) | n | Q ($\mu F$) | $R_s(\Omega)$ | $R_{ct}(G\Omega)$ | n | Q ($\mu F$) |
|  | 716.45 | 147.93 | 0.832 | 10.22 | 705.58 | 9.26 | 0.893 | 11.7 |
| | $\pm 74.59$ | $\pm 30.28$ | $\pm 0.037$ | $\pm 1.18$ | $\pm 45.68$ | $\pm 1.56$ | $\pm 0.016$ | $\pm 4.28$ |
| Stimulated channels | $R_s$ ($\Omega$) | $R_{ct}$ ($G\Omega$) | n | Q ($\mu F$) | $R_s(\Omega)$ | $R_{ct}(G\Omega)$ | n | Q ($\mu F$) |
| Electrode 1 | 702.2 | 135.3 | 0.854 | 10.98 | 728.7 | 6.41 | 0.906 | 21.13 |
| Electrode 2 | 660.6 | 162.5 | 0.857 | 11.56 | 719.8 | 9.15 | 0.907 | 14.78 |
| Electrode 3 | 648.1 | 164.7 | 0.828 | 10.67 | 724.3 | 11.13 | 0.888 | 19.35 |
| Electrode 4 | 636.7 | 185.4 | 0.843 | 10.58 | 617.8 | 10.7 | 0.910 | 14.94 |
| Electrode 5 | 725.8 | 103.2 | 0.748 | 9.61 | 690 | 5.17 | 0.901 | 8.767 |
| Electrode 6 | 689.5 | 157.4 | 0.834 | 8.34 | 677.8 | 6.63 | 0.871 | 8.73 |

Interestingly, the stimulated channel (E2C2) with increased *E_ac_* only presents minor damage (3.44 ± 0.19% in electrode area) at the electrode perimeter in optical micrographs (Fig. 3g and h, highlighted in red dot box). Likewise, models fit the Bode plots of E2C2 well for data from before and after Stim-Stab (Fig. 3j). Hence, the representative stimulated electrodes E1C2 and E2C2 with the most significant damage, optically remain functional with largely preserved electrochemical properties after the Stim-Stab test.

Thermoforming is used to shape *MouseFlex* electrodes to make intimate contact with the cervical vagus nerve. This procedure mechanically forms the electrode around a thin (127 µm) tungsten rod with a heat treatment in room-air to shape it. The tungsten mandrel imparts a radius of curvature as small as 65 µm, and therefore requires careful evaluation for damage to the electrodes. The process also exposes electrodes to temperatures near 260 °C in room air, which can impact the structure and electrochemical properties of the electrode stack and encapsulation. Fig. 4a presents the optical image of the integrated *MouseFlex* electrode, after thermoforming. The measured radius of curvature for the electrodes was 86 ± 12 µm. In comparison, a micro sling cuff electrode (°AirRay Cuff Sling Electrodes, CorTec, Freiburg, Germany) designed for very fine nerves exhibits the inner diameter of 200 µm, with 100 µm diameter electrodes also available. Noller *et al*. suggested that the inner diameter of cuff electrodes should be approximately 1.4 times larger than the outer diameter of the target nerve for rodent models, according to their experience and other published reports [29]. This ratio is potentially impacted by their mechanical characteristics, since these can impact tethering forces, constriction of the nerve, and trauma. Thermoformed electrodes are well-sized to make intimate contact with the mouse vagus nerve (r ≈ 50 µm). EIS Bode plots in Fig. 4b were collected before and after thermoforming. The circular markers represent the averaged spectra before and after thermoforming, with shaded error bars as standard deviation. The impedance magnitude and phase angle are statistically compared at each frequency to identify regions of significant changes. For frequencies upto 252 Hz, impedance magnitude |Z| from before and after thermoforming do not significantly change (*p* > 0.05). At middle frequencies (252 ~ 10^3^ Hz), the difference in |Z| after thermoforming becomes significant (*p* < 0.05), and at 10^3^ Hz the mean impedance of thermoformed electrodes is almost 57% higher than before the thermoforming process. The impedance spectra from before and after thermoforming is relatively constant at high frequency region (10^3^ ~ 10^5^ Hz), which is consistent with their near 0° phase angle and associated resistive character of electrodes in the access impedance regime. In contrast, the phase angle increases (becomes less negative) by a small but statistically significant (*p* < 0.05) amount from 1.5 Hz to 395 Hz. The phase angles then approach 0° and flatten at higher frequencies, which is consistent with resistive character associated with the access resistance.

**Fig. 4.**
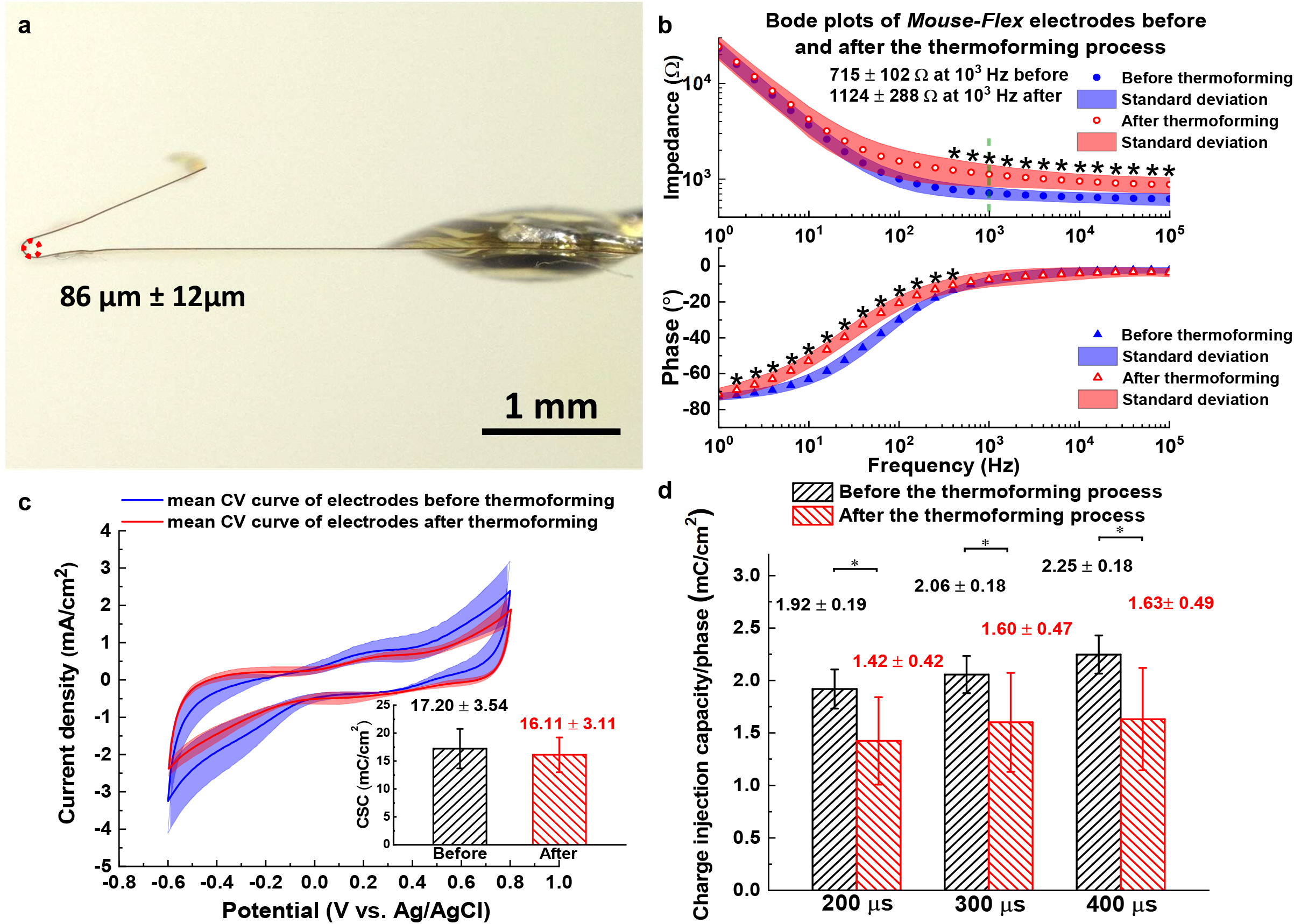
Thermoformed *MouseFlex* electrodes and its electrochemical properties. a) optical image of the thermoformed *MouseFlex* electrode with highlighted radius of 86 µm ± 12 µm, b) impedance spectra, c) charge storage capacity, and d) charge injection capacity of the *MouseFlex* electrodes.

The access impedance is a function of specific impedance for the PBS electrolyte, the surface area, and the spreading impedance between the working and counter electrodes. The small inner diameter of the cuff, the close proximity of the electrodes on the opposing, the lack of observed damage (e.g. delamination with thermoforming), controls for specific impedance of the electrolyte, and the fact that predominantly the access resistance dominated region of the spectra is impacted together suggest that the increase in impedance is the result of an increase in the spreading impedance.

The average CV curves of the *MouseFlex* electrodes before and after thermoforming (blue and red traces, respectively) and their associated standard deviations (shaded error bands) are presented in Fig. 4c. In agreement with previous studies [30–32], there are no strong redox peaks in CV curves of unthermoformed IrO_x_ electrodes. The CV curves are consistent indicating uniformity in the electrode quality associated with the microfabrication process. The average slope of the CV decreases after thermoforming which is consistent with a higher resistance. This is consistent with increased access impedance that also occurred with thermoforming. After thermoforming, no obvious redox peak was observed, suggesting the chemical state of the IrO_x_ layer did not significantly change due to thermoforming. Moreover, the CSC decreases from 17.20 ± 3.54 mC/cm^2^ to 16.11 ± 3.11 mC/cm^2^ after thermoforming, but is not statistically significant (*p* > 0.05). In contrast, the CIC was observed to statistical decrease (*p* < 0.05) for the pulse widths of 200, 300 and 400 µs (Fig. 4d). The mean CIC values at 200, 300 and 400 µs declines by 25.8%, 22.1% and 25.4%, respectively.

The efficacy of thermoformed *MouseFlex* electrodes was evaluated by stimulating the cervical vagus nerve of mouse models (n=8) and monitoring heart rate and respiration rate as biomarkers. Bradycardia and bradypnea or apnea, slowed heart rate or breathing rate, respectively, are two of the most commonly used biomarkers for VNS, putatively attributed to the Herring-Breuer reflex. They represent the most commonly used and reliable indicators of vagus nerve engagement used in bioelectronic medicine research [33,34].

A midline cervical incision made to expose the trachea is illustrated in Fig. 5a, and a higher magnification schematic diagram in Fig. 5b depicts the thermoformed *MouseFlex* electrode wrapped around the vagus nerve. The optical image (Fig. 5c) reveals intimate contact between the mouse vagus nerve and the thermoformed electrode. Fig. 5d and e present the percentage change in heart rate (∆HR) as a function of stimulation current for channel 1 and 2, respectively. The plots also include a vertical dashed line and shaded error bars representing the *in-vivo* charge injection capacity (mC/cm^2^) for the electrodes and its standard deviation. Stimulation with both channels 1 and 2 (stimulated in randomized order) shows similar trends for decreases in heart with increasing stimulation current. A roughly 20% decrease in heart rate is observed at 200 µA of current, and increasing stimulation current further decreases heart rate until the effect levels out near a 60% decrease.

**Fig. 5.**
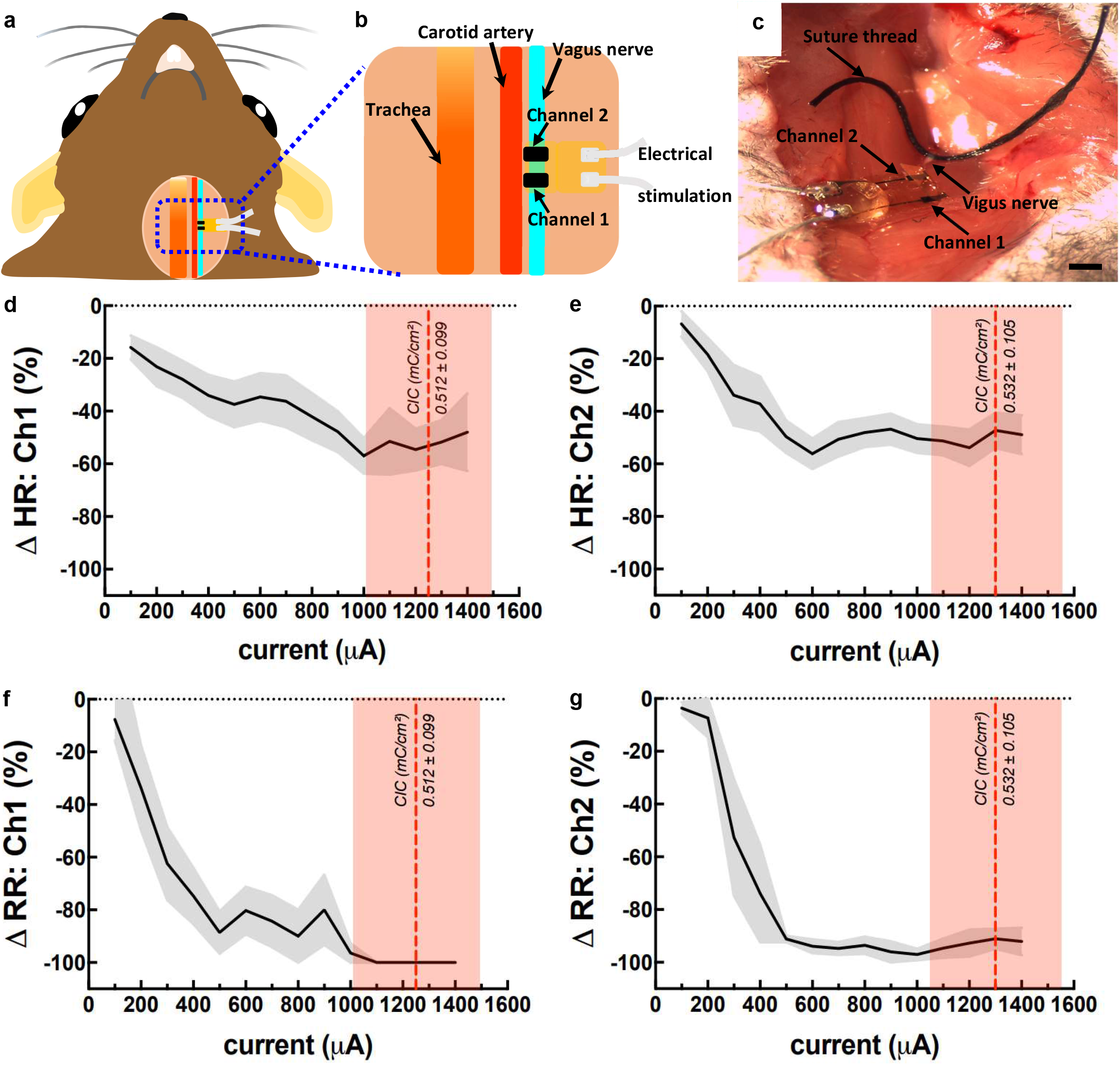
Acute VNS in mouse model and monitoring their heart and respiration rate as biomarkers for stimulation efficacy. a) schematic diagram of the acute VNS in the mouse model, b) zoom-in schematic diagram of the acute VNS in the mouse model, c) optical image of the acute VNS in the mouse model, d) mouse heart rate change as the function of stimulation current at channel 1, e) mouse heart rate change as the function of stimulation current at channel 2, f) mouse respiratory rate change as the function of stimulation current at channel 1, g) mouse respiratory rate change as the function of stimulation current at channel 2, scale bar =1 mm.

The threshold current to modulate the heart rate is commonly defined as resulting in a 5 ~ 10% drop in the heart rate [35]. Accordingly, the threshold current to modulate heart rate for both channel 1 and 2 is 100 or 200 µA. Differences were observed between the stimulation outcomes from each channel for the cohort, with channel 1 having a stronger onset effect at 100 µA, but a shallower slope for the relationship between degree of bradycardia and stimulation compared to channel 2. The *in-vivo* CICs were also quantified by collecting and analyzing the nerve stimulation voltage transients and characterizing the maximum current where *E*_mc_ is within the water window (> −0.6 V). The so-called *water window* is generally accepted as a boundary for safe nerve stimulation to avoid gas bubble formation, changes in *p*H, or damage to the electrodes resulting in poor outcomes for the nerve and/or electrode. The CICs determined for channel 1 and 2 are 0.512 ± 0.099 mC/cm^2^ (at 1250 ± 243 µA) and 0.532 ± 0.105 mC/cm^2^ (at 1300 ± 256 µA), respectively. The small difference between CIC values is not statistically significant (*p* > 0.05). These values are roughly 32% of the CIC values for the thermoformed electrodes measured in PBS at 37°C, which are the most directly comparable electrodes for this evaluation. *In-vivo* CIC values are much less commonly measured, and are critical to truly evaluating safety and efficacy for electrodes, and are very often considerably lower than values measured in saline.

Fig. 5f and g presents the change in respiration rate (∆RR(%)) compared to baseline as a function of stimulation current for channel 1 and 2, respectively. Similar to the trends observed for ∆HR, the ∆RR quickly decreases towards 100% (apnea) for both channel 1 and channel 2, with increasing current. The threshold current to modulate RR is defined as the stimulation current generating a 5 ~ 10% change (drop) in the RR. Channel 1 and 2 exhibit threshold currents of 100 µA and 200 µA, respectively. The ∆RR for both electrodes exceeds −90% at 500 µA, which is consistent with the near or complete apnea visually observed during the 7 second stimulation epochs. The effect of stimulation on the RR is larger than that on HR, which is consistent with observations in the literature regarding VNS [36].

The electrochemical properties of *MouseFlex* electrodes (EIS, CSC, and CIC) measured before, during and after surgical placement of the electrodes were compared to evaluate the effects of surgical handling and also how their *in-vivo* and *in-vitro* characteristics compare. Fig. 6a presents impedance spectra from before, during, and after placement on the mouse vagus nerve. The |Z| while placed on the mouse vagus nerve is significantly higher and has a larger standard deviation (shaded error bands), compared to both before and after placement on the nerve at frequencies > 10 Hz. Non-specific protein binding, differences in tissue spreading impedance compared to saline, and difference in the 3-electrodge geometry are potential factors contributing to the higher impedance. Surprisingly, the impedance after extracting and cleaning the electrodes when measured in saline does not significantly change compared to before placement for frequencies above 100 Hz, and slightly decreased at low frequencies (~10 Hz). The latter effect suggests that the IrO_x_ electrode was slightly activated during *in-vivo* use. Overall this strongly suggests that neither the surgical handling or cleaning process degrade the electrode impedance characteristics. The impedance phase angle (Fig. 6b) after *in-vivo* use is also changes little, except for a slightly steeper slope in the transition between the capacitive (low frequency) and resistive (higher frequency) regimes. There was also a slight offset of this transition towards lower frequencies. In comparison, phase angles for the electrodes on the vagus nerve had predominantly resistive character for frequencies > 100 Hz, associated access resistance. This in conjunction with the higher access impedance in this region could result from several mechanisms, which requires significant further experimentation to understand. The slope towards lower phase angles at low frequencies is less steep than those measured in saline, suggesting a perturbation to the electrochemical interface. The mechanisms for this could be factors such as non-specific protein binding, cellular adhesion, or other factors adsorbing or adhering to the electrode and changing its electrochemical properties.

**Fig. 6.**
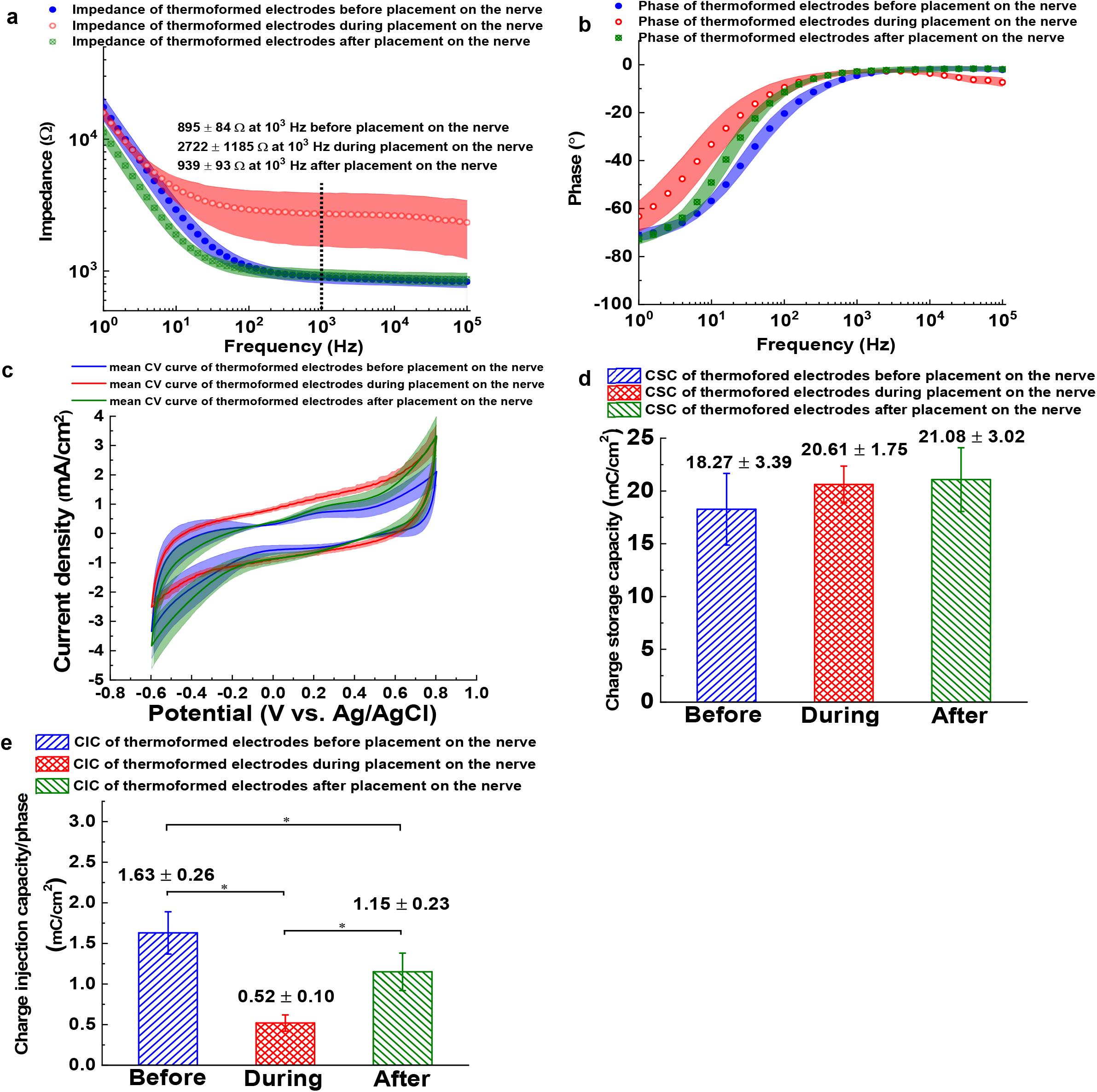
*In-vivo* electrochemical properties of thermoformed *MouseFlex* electrodes before, during and after placement on the nerve. a) impedance spectra of thermoformed *MouseFlex* electrodes before, during and after placement on the nerve, b) phase spectra of thermoformed *MouseFlex* electrodes before, during and after placement on the nerve, c) mean CV curves spectra of thermoformed *MouseFlex* electrodes before, during and after placement on the nerve, d) CSC of thermoformed *MouseFlex* electrodes before, during and after placement on the nerve, e) CIC of thermoformed *MouseFlex* electrodes before, during and after placement on the nerve.

The CV curves of electrodes before, during and after placement are presented in Fig. 6c. No strong redox peaks were observed in any of the CV curves though more features are clearly presented from samples measured in saline before and after placement on the vagus. The *in-vivo* CV curves were more consistent with narrow standard deviation and take on a more ideally capacitive character with the only electrochemical feature being reduction and oxidation of water (electrolysis). Furthermore, despite slight morphological differences between the curves, there were no statistical difference in any of the CSC values (Fig. 6d). This lack of change in CSC during nerve placement is surprising, given the significant changes in impedance and CIC (described in the following paragraph). The stability of the CSC during and after nerve placement further supports that the electrodes are sufficiently robust to tolerate thermoforming and surgical handling.

We also evaluated the CIC before, during, and after placement on the vagus nerve, which are presented in Fig. 6e. The CIC at 30 Hz with the pulse-width of 260 µs before placement on the nerve is very similar to the higher frequency measurements in saline presented earlier (1 kHz, 300µs, Fig. 4g, *p*>0.05). This suggests that the pulse rate does not significantly impact the CIC within the tested frequencies. Interestingly, the CIC of electrodes during placement on mouse vagus nerves very significantly (*p* < 0.05) decreases to a value only one fifth of that before placement. However, after removal and a cleaning process, the CIC of electrodes returns to 70.6% of original values when measured in PBS on the benchtop, which is also a significant increase compared to the *in-vivo* CIC (*p* < 0.05), but surprisingly not a significant change compared to the value before placement (*p* > 0.05). Unreported experiments with *Flex* electrodes and previously reported results from Utah arrays also found that cleaning with Enzol did not effect the electrochemical properties of electrodes [37].

## 4. Discussion

Electrode stability under harsh electrical stimulation is a very useful engineering tool for optimizing the stability of electrodes materials and devices, as they are developed for *in-vivo* use and clinical translation. The stability of flexible IrO_x_ electrodes, particularly after thermoforming, has not been systematically investigated. Additionally, *in-vivo* electrochemical changes that occur and their implications on device safety and efficacy require further attention. A recent study investigated stability of SIROF deposited on a Utah electrode array, which was deposited with gas composition incorporating water vapor in the plasma. These electrodes had small changes in CV curves and slightly reduced 1 kHz-impedance after 10^9^ cycles of stimulation at 0.4 mC/cm^2^/phase [30]. In this research, we investigated the stimulation stability of IrO_x_ at higher charge density (1.51 mC/cm^2^/phase) for PI substrates, along with careful evaluation on the effects of thermoforming and electrochemical properties before, during and after surgical placement along with optical microscopy.

All electrodes remain electrically and electrochemically (impedance, CSC and CIC) functional after 10^9^ stimulation pulses at 1.51 mC/cm^2^/phase, although 2 of 6 electrodes had observed discolorations in optical micrographs and relatively large variation of voltage transient waveforms suggesting degradation processes for the electrodes are occurring. The observed degradation suggests the electrodes are possibly in early failure distribution due to 10^9^ pulsing stimulation at 1.51 mC/cm^2^/phase. The achieved stimulation lifetimes suggest the electrodes are compatible with the *in-vivo* stimulation conditions involved with chronic animal studies.

Thermoforming *MouseFlex* electrodes to interface small peripheral nerves is a thermal mechanical process to reshape the planar IrO_x_ electrodes into 3D structures. The capability to withstand the thermal mechanical process at high temperature without serious degradation on electrochemical properties, mechanical properties, lifetime, and biocompatibility is the prerequisite for the planar flexible electrode in translational studies. It was reported that thermoforming at 450 °C did not have influence on chemical composition or chemical resistance of a PI film (Kapton® HN, Dupont Inc., Wilmington, DE), but the thermoformed PI exhibited higher elasticity than pristine PI [38]. As evidenced by little difference in CV curves, sensing performance of a PI gold electrode encapsulated by polyurethane remained unchanged after a low-temperature thermal moulding (<150 °C) [39]. Lee *et al* developed a thermoforming process at 230 °C to wrap flexible PI (PI 2611) cuff electrodes around 800 µm-diameter rat sciatic nerve [40]. Compared to these reports, the thermoforming process in this study for the *MouseFlex* electrodes (PI 2611) is more harsher to likely induce more inner stress and damage to IrO_x_ layer due to smaller wrapped diameter (~127 µm) and higher temperature (~260 °C).

Moreover, from the perspective of neural interface, systematic electrochemical characterization was thoroughly performed in this study to investigate the impact of the thermoforming on the electrochemical properties of the *MouseFlex* electrodes. Although the mean impedance of electrodes at 1 kHz significantly increases by 57%, no statistical difference was found in CSC after thermoforming. The CIC of electrodes decreased by nearly 30% was observed after thermoforming. Potential mechanisms are damage to the electrode metallization or increased polarization due to constricted volume of stimulated electrolyte within the cuff. For *MouseFlex* electrodes, CIC is more sensitive than CSC as a criteria to evaluate changes in the electrode from the thermoforming process, and understanding the mechanisms for these changes remains of interest for future work.

The stimulation current for mouse vagus nerve and peripheral nerves, depends on the character of the electrode-tissue interface, electrode architecture, as well as biotic factors such as tissue response and surgical approach. In a seminal study of mouse VNS, the stimulation current threshold of a custom bipolar cuff electrode (Microprobes for Life Science, Gaitherburg, MD) was 500 µA to activate inflammatory reflex [41]. Similarly, stimulation current of 300 µA dramatically supressed mice plasma amylase and lipase concentration [42]. Mice VNS at 50 µA using a bipolar silver Cooner wire electrode elicited significant 10% ~ 15% reduction in HR. While with the implantation up to 1 week, the stimulation current threshold to lead to 5% ~ 15% drop in HR was reported to increase to approximately 400 µA for a Micro-leads cuff electrode [43]. A low stimulation current threshold for vagus nerve resulting in physiological responses (e.g. decrease in HR and RR) is desirable to minimize irreversible electrochemical reactions and tissue damage to achieve safe VNS. The microfabricated *MouseFlex* electrodes demonstrated the relatively low stimulation current threshold to modulate HR (~100 µA for both channel 1 and 2) and RR (~100 µA for channel 1, ~200 µA for channel 2), validating clear efficacy and suggesting excellent charge delivery characteristics.

The limits for safe stimulation of the mouse vagus nerve, particularly for chronic studies, requires significant further investigation [44]. Ultimately, histological tissue analysis is required to determine safe stimulation paradigms for a given electrode and study endpoint, but some guidelines are available in the literature including a review paper [45]. Large stimulation currents can cause nerve damage, resulting from gas evolution due to potentials beyond the water window, formation of toxic by-products, excitotoxicity, electroporation, and thermal effects. The water window is widely regarded as an upper bound for safe stimulation, therefore we evaluated our ability to evoke physiological responses at threshold currents lower than those resulting in voltages exceeding the water window. Although we did not perform a histological investigation, the lack of acute damage to the nerve is further supported by our ability to evoke responses from both the second electrode used for stimulation (channel 1 and channel 2 in randomize order). The combination of 100 and 200 µA threshold currents and water window currents (1250 ± 243 mA for channel 1, and 1330 ± 256 mA for channel 2) strongly suggest safe and effective stimulation can be achieved within our paradigm.

Compared to bench CIC measurements in saline solution, a significant decline in *in-vivo* CIC was observed from our acute study. This is also consistent with chronic electrode implantation. The mean *in-vivo* CIC of 11.5% to 30.8% of the *in-vitro* CIC, when Pt electrodes were acutely or chronically implanted into suprachoroidal space, and values within this range were also found for cortical electrodes [46]. Likewise, the mean *in-vivo* CIC of TiN coated PtIr electrodes were found to be 37.1% of the mean *in-vitro* CIC, when implanted in the rat motor cortex [47]. Utah electrode arrays with IrO_x_ tip metallization exhibited similar behaviour in cat cortex, presenting *in-vivo* CIC values that decreased by a factor of 2 to 3 [48]. The *in-vivo* CIC of PI based microelectrodes using IrO_x_ electrode sites have been evaluated by the Stieglitz lab with values of 60 nC/ph (nominally 1.19 mC/cm^2^/ph) for 80 µm diameter electrode sites [13]. Smaller electrodes sites are widely reported to have higher CIC values, and these electrodes are 25× smaller than the *MouseFlex* sites. The *in-vivo* CIC of flexible IrO_x_ electrodes thermoformed and placed on mouse vagus nerve has not been evaluated for VNS. The mean *in-vivo* CIC values (0.512 ± 0.099 mC/cm^2^ for channel 1, and 0.532 ± 0.105 mC/cm^2^ for channel 2) which compares favorably with state-of-the-art values for IrO_x_-based electrodes. Importantly, we are able to achieve robust vagus nerve stimulation well below the CIC, which supports the safe and efficacious capability for *MouseFlex* electrodes.

Conducting polymers are another very promising electrode material with low impedance, high CSC, excellent biocompatibility and chemical stability, and have been widely reported for neural signal recording [49]. A variety of methods such as electrografting P(EDOT-NH2) are being evaluated as adhesion-promoting layer, which is proposed to form the covalent bonds between organic species and metal or metal oxide substrates [50]. To our knowledge, the ability of PEDOT to withstand the thermal and mechanical rigorous of thermoforming for small peripheral nerves, and the ability of these coatings to handle high stimulation current densities (near the water window) have not been studied. Performance of PEDOT coatings on IrO_x_ in recent studies [51], which are potentially exciting future studies.

Flexible polymer electrodes and thin-film electrode metallization are potentially more vulnerable to damage during surgerical use and handling compared to traditional Pt foil electrodes in silicone used for deep brain stimulation and cochlear electrodes. This makes evaluating the electrodes ability to tolerate surgical handling and use a critical part of testing. Stieglitz *et al*. evaluated PI-based transverse intrafascicular multichannel electrodes (TIMEs) explanted after use to stimulate the median nerve of a human subject, and leg nerves in animal models [52,53]. Fan *et al*. compared the impedance of sputtered porous Pt electrodes on SU-8 substrate, before brain implantation and after removal and subsequently soaking in distilled water [54]. We studied the tolerance of our electrodes for surgical handling by carefully recovering the placed electrodes, soaking them in enzymatic detergent solution overnight and thorough rinsing and irrigation with DI water. This removed the vast majority of tissue, proteins and organic matter on *MouseFlex* electrodes. The impedance (magnitude and phase), CSC, and CIC were then measured and compared. The impedance and CSC, did not significantly change after surgical handling and cleaning process. Similar to the thermoforming study, CIC is more sensitive to the effects of surgical handling, and experience a roughly 30% decrease in CIC. This relatively high degree of electrochemical stability for the electrodes in combination with their ability to evoke robust physiological responses with stimulation suggests they are well stuited for acute use, and is that chronic evaluation with suitably adapted electrodes is warranted.

## 5. Conclusions

In this study, we present details regarding bench and *in-vivo* evaluation of recently developed electrodes (*MouseFlex*). These PI-based electrodes are microfabricated and use IrO_x_ for electrode sites, and are thermoformed into a cuff geometry for the small-diameter mouse vagus nerve (r ≈ 50 µm). The MEMS approach enables scalable and repeatable fabrication. The IrO_x_ electrodes exhibit low impedance (762 ± 96 Ω at 1 kHz), relatively high CSC (13.92 ± 3.01 mC/cm^2^) and CIC (2.23 ± 0.66 mC/cm^2^ at 400 ms). The stability of the IrO_x_ electrode sites was evaluated by a Stim-Stab test applying 10^9^ stimulation pulses at a charge density of 1.51 mC/cm^2^/phase. Although this is one of the harshest stimulation protocols reported, all electrodes remained functional, as determined by a comprehensive evaluation including electrochemical properties (impedance (with ECM), CSC and CIC), real-time VT, *E_mc_* and *E_ac_* curves, and optical images. A thermoforming process was developed to shape the 2D *MouseFlex* electrodes into a 3D structure, and create an intimate neural interface with vagus nerve. The thermoforming process had a mild negative effect on the electrochemical properties of the electrodes, which could result from constriction of the spreading resistance through the electrolyte or damage to the electrode metallization. The stimulation efficacy of the thermoformed *MouseFlex* electrodes was demonstrated by decreasing the heart and respiration rates at a threshold current of 100 µA and 200 µA, respectively. Importantly, the *in-vivo* CIC of the thermoformed *MouseFlex* electrodes placed on vagus nerves were measured to be 0.512 ± 0.099 mC/cm^2^, corresponding to an current amplitude of 1250 ± 243 µA as the limit for safe mouse VNS. The electrochemical properties were assessed before, during and after *in-vivo* use, enabling a careful analysis of the impacts of these on the electrochemical properties. Finally, thermoformed *MouseFlex* electrodes are sufficiently robust to tolerate the surgical handling, as supported by comparable and reversible electrochemical properties before and after electrode placement on the mouse vagus nerve. These flexible electrodes for interfacing with small-diameter nerves should enable additional mechanistic studies within the fields of bioelectronic medicine and neuromodulation.

## Acknowledgement

This work was partially supported by industry contracts with General Electric and United Therapeutics, and with internal funding from West Virginia University and Northwell Health. The *MouseFlex* electrodes were fabricated at the Utah Nanofabrication Lab.

## Supplementary Materials

**Figure S1**. Representative VT waveform of a Mouse Flex electrode in response to a charge-balanced, biophasic, cathodic-leading, symmetrical current pulsing.

**Figure S2**. The stimulation paradigm used for the Stim-Stab test.

**Figure S3**. The thermoforming process to reshape the *Mouse Flex* electrode around the tungsten rod. a) heat the folded Mouse Flex electrode perpendicularly, b) heat the folded *Mouse Flex* electrode horizontally.

**Figure S4**. a) Real-time voltage transient (VT) waveform of 1 kΩ resistor at 8 mA, b) *E_ac_* and *E_mc_* curves of 1 kΩ resistor stimulated at 8 mA.

**Figure S5**. High-magnification optical images of a representative unstimulated channel (a) before and (b) after the Stim-stab test.

**Fig. S1.**
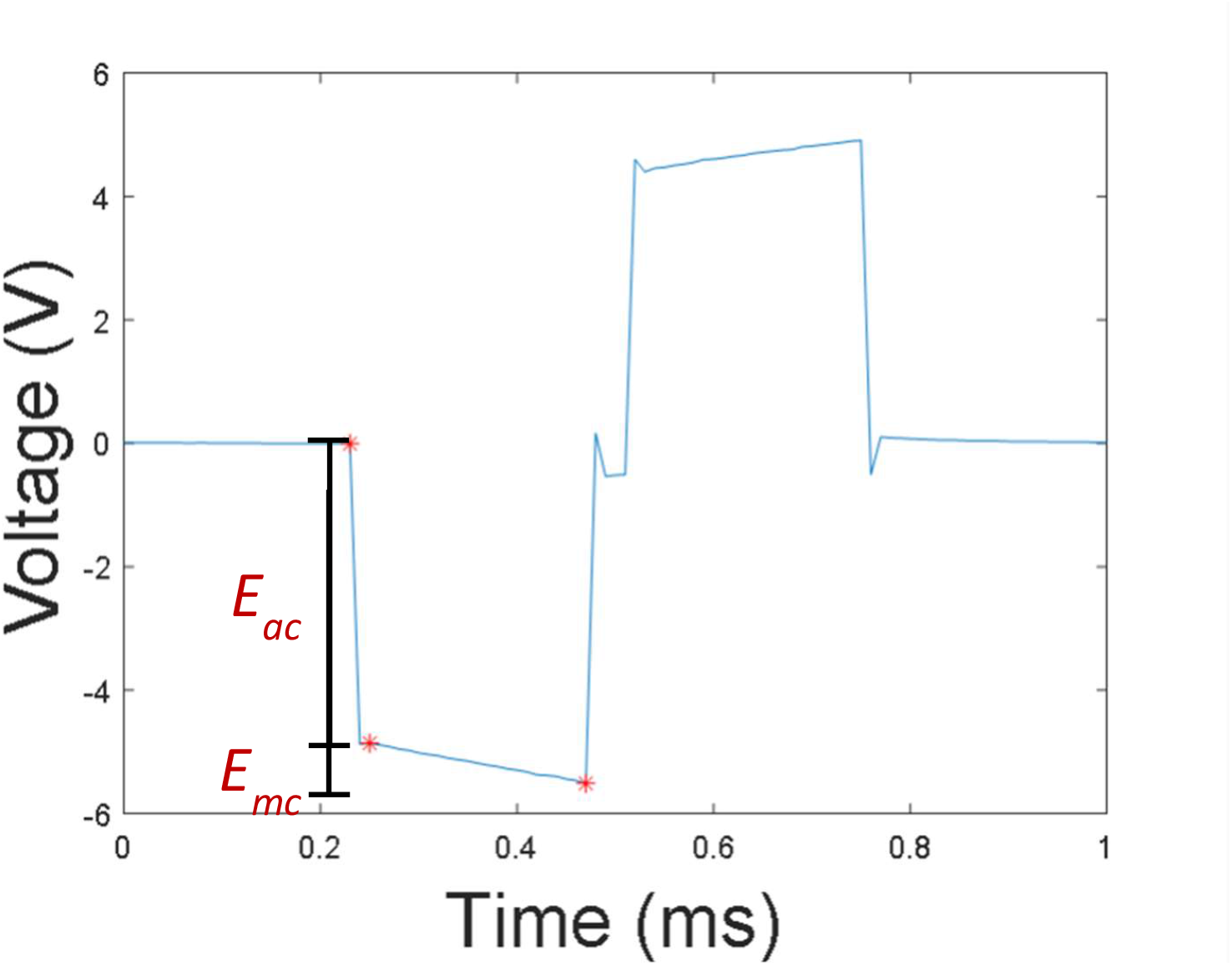
Representative VT waveform of a *MouseFlex* electrode in response to a charge-balanced, biophasic, cathodic-leading, symmetrical current pulsing.

**Fig. S2.**
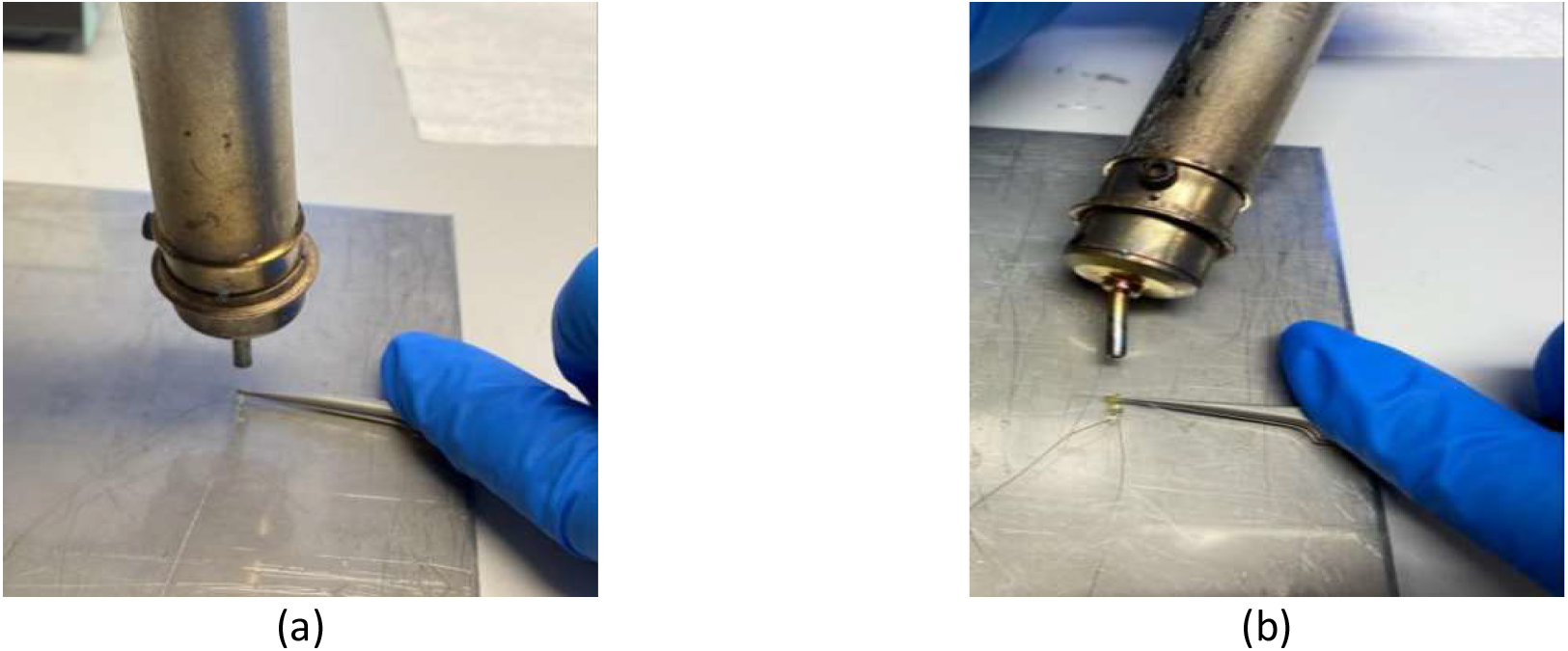
The thermoforming process to reshape the *MouseFlex* electrode around the tungsten rod. a) heat the folded *MouseFlex* electrode perpendicularly, b) heat the folded *MouseFlex* electrode horizontally.

**Fig. S3.**
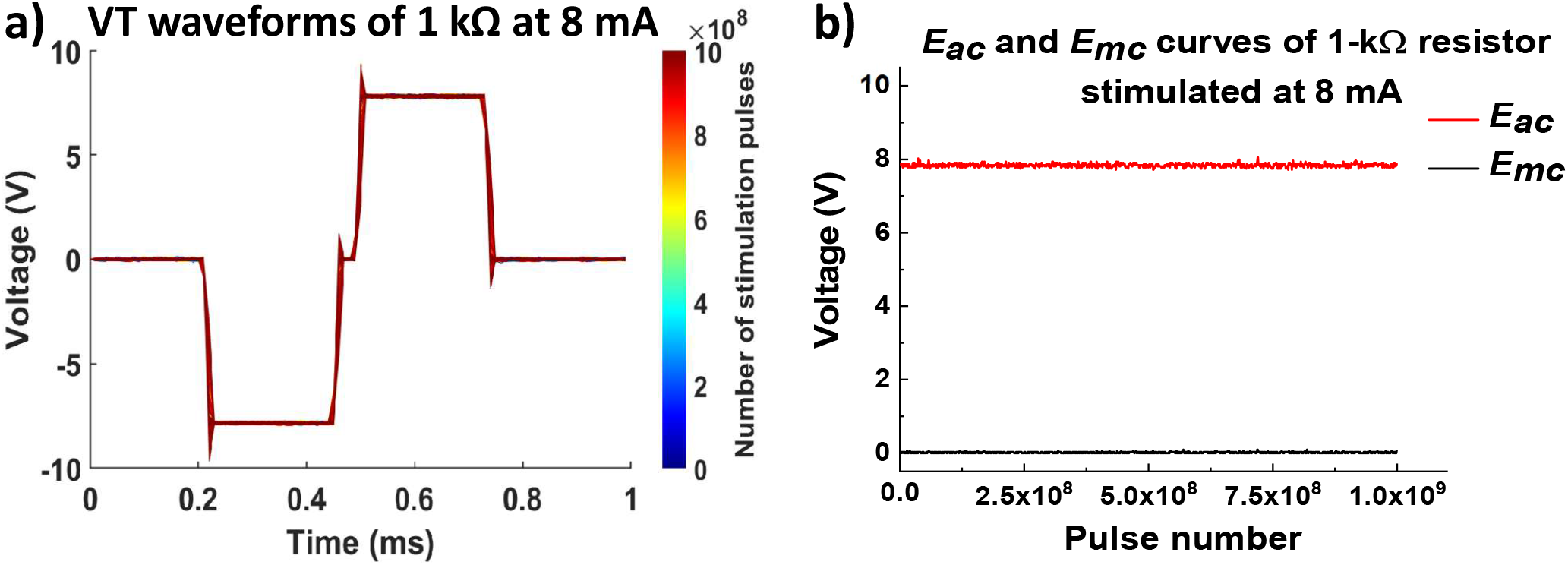
a) VT waveforms of 1 kΩ at 8 mA as the function of the stimulation cycle, b) *E_ac_* and *E_mc_* curves of 1-kΩ resistor stimulated at 8 mA.

**Fig. S4.**
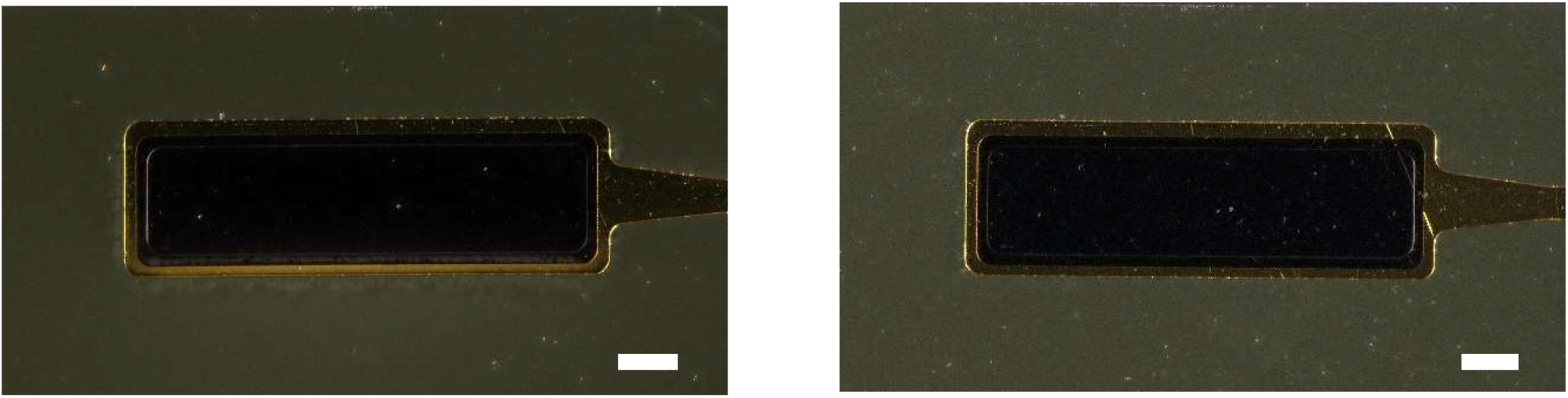
High-magnification optical images of a representative unstimulated channel (a) before and (b) after the Stim-Stab test, showing no damage occurred for unstimulated channel after the test (scale = 100 µm).

**Fig. S5.**
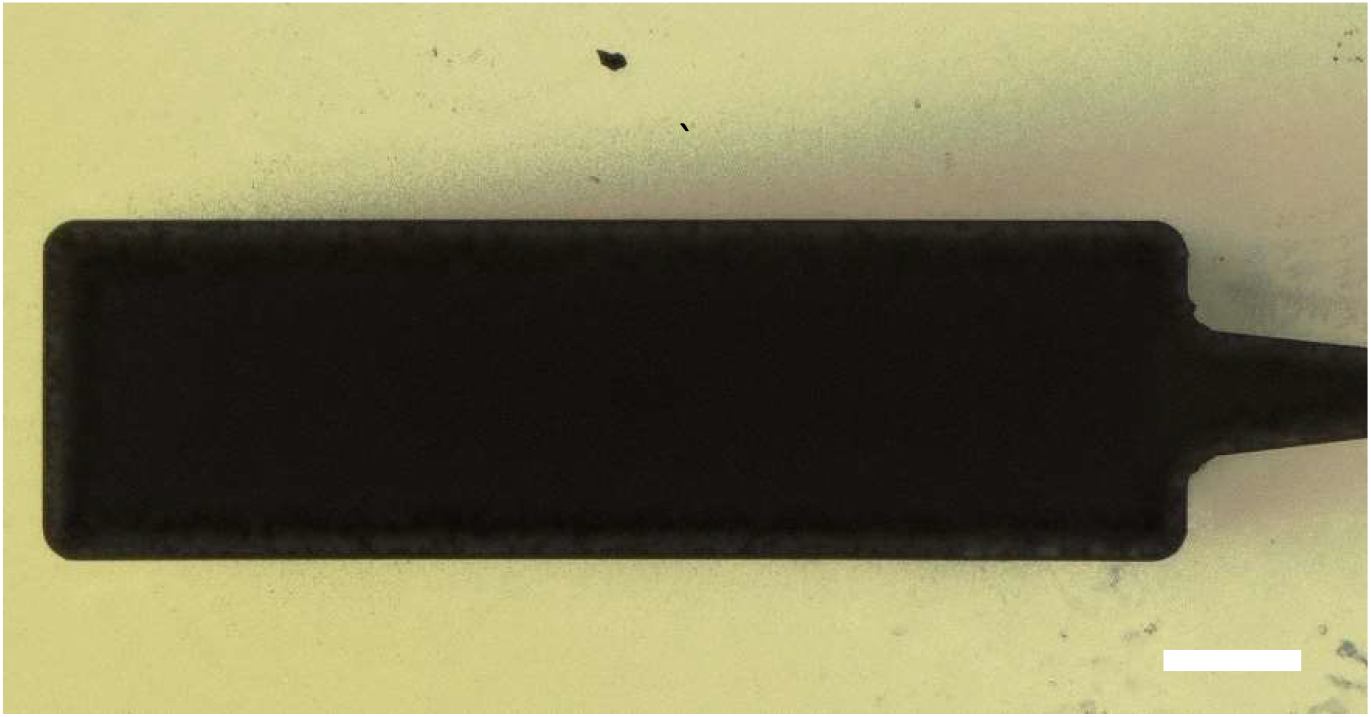
Transmitted optical image of the stimulated channel E1C1, showing 1) no stacked layer delamination from the electrode site where is opaque, 2) the discoloration area is not as transparent as PI layer, suggesting a metal layer formed at the interface of PI layers or within them (scale = 100 µm).

