## Supplemental materials for "Flexible IrO_x_ Neural Electrode for Mouse Vagus Nerve Stimulation"

### Supplementary Materials

**Figure S1.** Representative VT waveform of a Mouse Flex electrode in response to a charge-balanced, biophasic, cathodic-leading, symmetrical current pulsing.

**Figure S2**. The stimulation paradigm used for the Stim-Stab test.

**Figure S3.** The thermoforming process to reshape the *Mouse Flex* electrode around the tungsten rod. a) heat the folded Mouse Flex electrode perpendicularly, b) heat the folded *Mouse Flex* electrode horizontally.

**Figure S4.** a) Real-time voltage transient (VT) waveform of 1 kΩ resistor at 8 mA, b) $E_{ac}$ and $E_{mc}$ curves of 1 kΩ resistor stimulated at 8 mA.

**Figure S5.** High-magnification optical images of a representative unstimulated channel (a) before and (b) after the Stim-stab test.

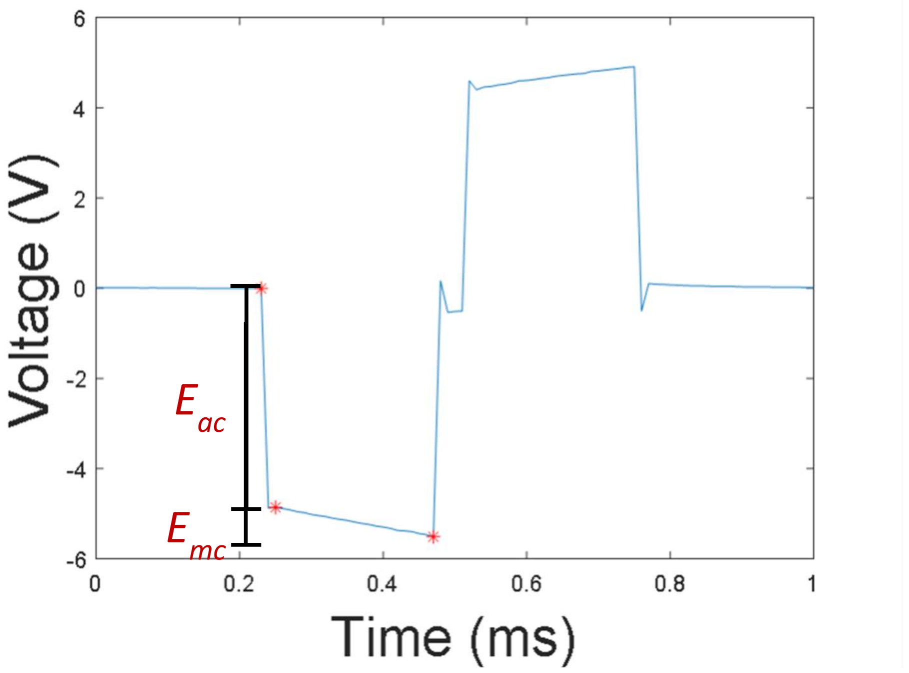


**Fig. S1.** Representative VT waveform of a *MouseFlex* electrode in response to a charge-balanced, biophasic, cathodic-leading, symmetrical current pulsing.

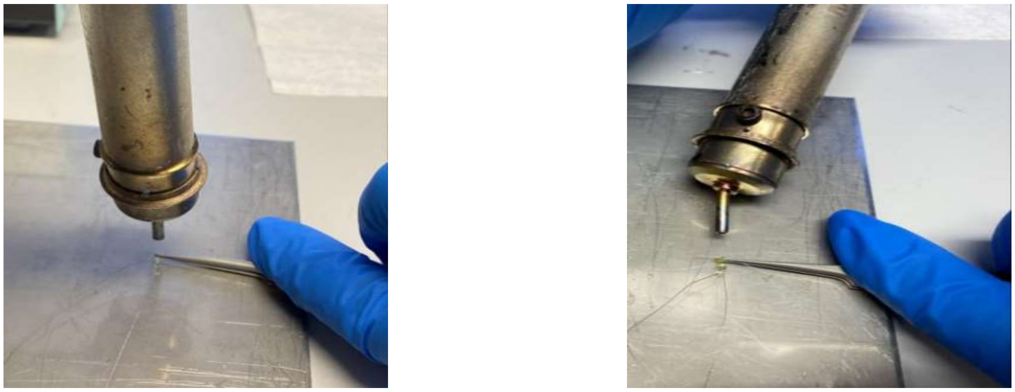

(a) (b)

**Fig. S2.** The thermoforming process to reshape the *MouseFlex* electrode around the tungsten rod. a) heat the folded *MouseFlex* electrode perpendicularly, b) heat the folded *MouseFlex* electrode horizontally.

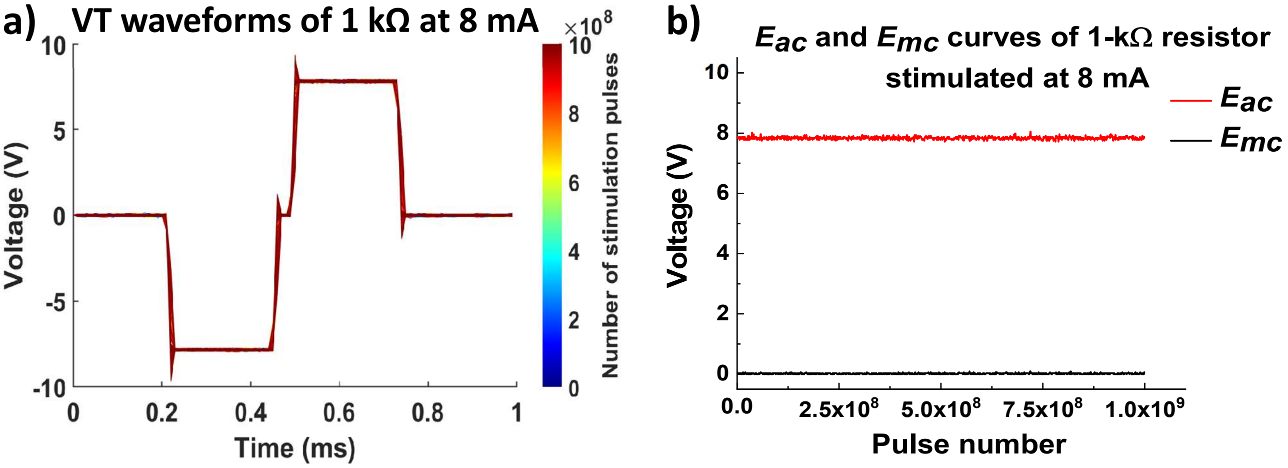


**Fig. S3.** a) VT waveforms of 1 kΩ at 8 mA as the function of the stimulation cycle, b) $E_{ac}$ and $E_{mc}$ curves of 1-kΩ resistor stimulated at 8 mA.

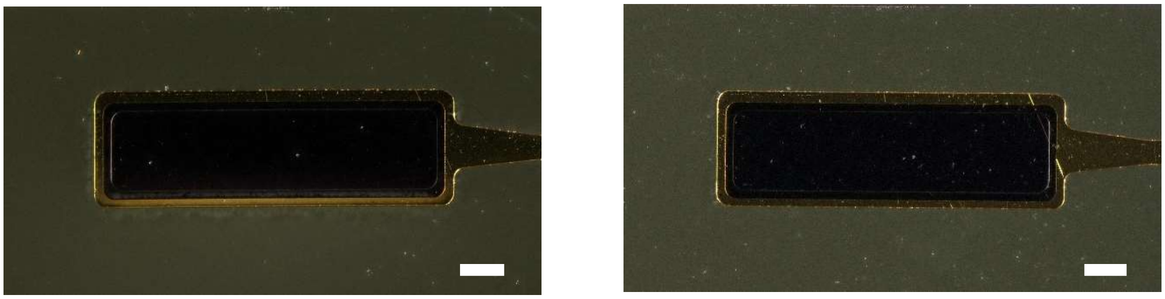

**Fig. S4.** High-magnification optical images of a representative unstimulated channel (a) before and (b) after the Stim-Stab test, showing no damage occurred for unstimulated channel after the test (scale = 100 μm).

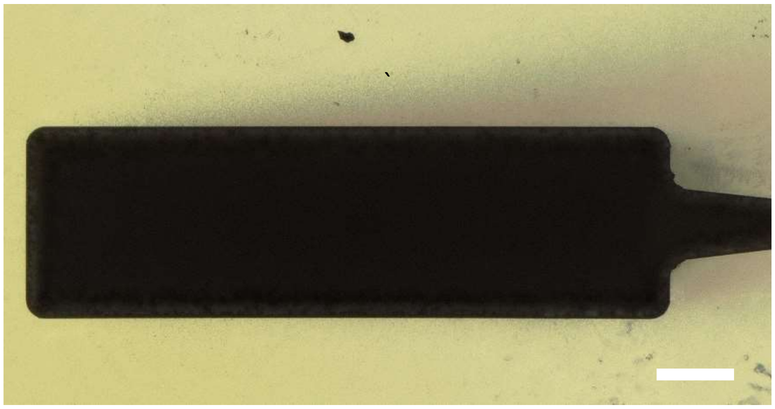

**Fig. S5.** Transmitted optical image of the stimulated channel E1C1, showing 1) no stacked layer delamination from the electrode site where is opaque, 2) the discoloration area is not as transparent as PI layer, suggesting a metal layer formed at the interface of PI layers or within them (scale = 100 μm).
